# A deep learning system can accurately classify primary and metastatic cancers based on patterns of passenger mutations

**DOI:** 10.1101/214494

**Authors:** Wei Jiao, Gurnit Atwal, Paz Polak, Rosa Karlic, Edwin Cuppen, Alexandra Danyi, Jeroen de Ridder, Carla van Herpen, Martijn P. Lolkema, Neeltje Steeghs, Gad Getz, Quaid Morris, Lincoln D. Stein, for the PCAWG Pathology and Clinical Correlates Working Group and the ICGC/TCGA Pan-cancer Analysis of Whole Genomes Network

**Author notes:** These authors contributed equally to the work.

## Abstract

In cancer, the primary tumour’s organ of origin and histopathology are the strongest determinants of its clinical behaviour, but in 3% of the time a cancer patient presents with metastatic tumour and no obvious primary. Challenges also arise when distinguishing a metastatic recurrence of a previously treated cancer from the emergence of a new one. Here we train a deep learning classifier to predict cancer type based on patterns of somatic passenger mutations detected in whole genome sequencing (WGS) of 2606 tumours representing 24 common cancer types. Our classifier achieves an accuracy of 91% on held-out tumor samples and 82% and 85% respectively on independent primary and metastatic samples, roughly double the accuracy of trained pathologists when presented with a metastatic tumour without knowledge of the primary. Surprisingly, adding information on driver mutations reduced classifier accuracy. Our results have immediate clinical applicability, underscoring how patterns of somatic passenger mutations encode the state of the cell of origin, and can inform future strategies to detect the source of cell-free circulating tumour DNA.

## Introduction

Human cancers are distinguished by their anatomic organ of origin and their histopathology. For example, lung squamous cell carcinoma originates in the lung and has a histology similar to the normal squamous epithelium that lines bronchi and bronchioles. Together these two criteria, which jointly reflect the tumour’s cell of origin, are the single major predictor of the natural history of the disease, including the age at which the tumour manifests, its factors, growth rate, pattern of invasion and metastasis, response to therapy, and overall prognosis. A tumour’s type is generally determined by a histopathologist who examines microscopic sections of the tumour, using non-specific stains, occasionally supplemented with protein-specific immunohistochemistry. However an increasing number of tumour types are subclassified using molecular markers that distinguish among subtypes with clinically distinct features.

Based on recent large-scale exome and genome sequencing studies we now know that major tumour types present dramatically different patterns of somatic mutation.^1-4^ For example, ovarian cancers are distinguished by a high rate of genomic rearrangements,^5^ chronic myelogenous leukemias (CML) carry a nearly pathognomonic structural variation involving a t(9;22) translocation leading to a BCR-ABL fusion transcript,^6^ melanomas have high rates of C>T and G>A transition mutations due to UV damage,^7^ and pancreatic ductal adenocarcinomas have near-universal activating mutations in the KRAS gene.^8^ Recent work has pointed to a strong correlation between the regional somatic mutation rate and chromatin accessibility as measured by DNase I sensitivity and histone mark^9^, and has suggested that the cell of origin can be inferred from regional mutation counts^10^.

This paper asks whether we can use machine learning techniques to accurately determine tumour organ of origin and histology using the patterns of somatic mutation identified by whole genome DNA sequencing. One motivation of this effort was to demonstrate the feasibility of a next-generation sequencing (NGS) based diagnostic tool for tumour type identification. Due to its stability, DNA is particularly easy to recover from fresh and historical tumour samples; furthermore, because mutations accumulate in DNA, they form a historic record of tumour evolution unaffected by the local, metastatic environment. Studies have shown that site-directed therapy based on the tumour’s cell of origin is more effective than broad-spectrum chemotherapy;^11^ however it is not always straightforward to determine the origin of a metastatic tumour. In the most extreme case, a pathologist may be presented with the challenge of determining the source of a poorly differentiated metastatic cancer when multiple imaging studies have failed to identify the primary (“cancer of unknown primary,” CUPS).^12^ A related challenge occurs when a patient has a past history of successfully treated cancer, and the pathologist is called upon to distinguish between a late recurrence of the disease versus a new cancer.

In current practice, pathologists use histological criteria assisted by immunohistochemical stains to determine such tumours’ histological type and site of origin^13^, but this process can be complex and time-consuming, and some tumours are so poorly differentiated that they no longer express the cell-type specific proteins needed for unambiguous immunohistochemical classification. Here we explored whether a simple DNA-based sequencing and analysis protocol for tumour type determination would be a useful adjunct to existing histopathological techniques.

A complementary motivation of this study is assessing the predictive power of various types of DNA mutations for classifying cancer type. As such, we tested the predictive accuracy of three broad categories of mutational feature: (1) the regional distribution of somatic passenger mutations, which bear the traces of the current and historical epigenetic state of the tissue of origin; (2) the distribution of somatic mutation types, which reflect environmental and genetic exposures of the cell of origin; and (3) the driver genes and pathways that are altered in the tumour. Unexpectedly, we found that passenger mutation regional distribution and mutation type are sufficient to discriminate among tumour types with a high degree of accuracy, while driver genes and pathways contribute provide no improvement to the classifier and perform badly at classifying cancer type when used on their own.

## Results

Using the Pan-cancer Analysis of Whole Genomes (PCAWG) data set,^4^ we built a series of tumour-type classifiers using individual sequence-based features and combinations of features. The best performing classifier was validated against an independent set of tumour genomes to determine overall predictive accuracy, and then tested against a series of metastatic tumors from known primaries to determine the accuracy of predicting the primary from a metastasis. We also examined patterns of misclassification errors to identify cases in which different tumour types share similar biology.

### Tumour Types

The full PCAWG data set consists of tumours from 2778 donors comprising 34 main histopathological tumour types, uniformly analysed using the same computational pipeline for quality control filtering, alignment, and somatic mutation calling. However, the PCAWG tumour types are unevenly represented, and several have inadequate numbers of specimens to adequately train and test a classifier. We chose a minimum cutoff of 35 donors per tumour type. In a small number of cases, the same donor contributed both primary and metastatic tumour specimens to the PCAWG data set. In these cases we used only the primary tumor for training and evaluation, except for the case of the small cohort of myeloproliferative neoplasms (Myeloid-MPN; N=55 samples), for which multiple primary samples were available. In this case, we used up to two samples per donor and partitioned the training and testing sets to avoid having the same donor appear more than once in any training/testing set trial. The resulting training set consisted of 2436 tumours spanning 24 major types (Table 1 and Supplementary Table 1).

**Table 1:** Distribution of tumour types in the PCAWG training and test data sets

| Abbreviation | Organ system | Tumor Type | Tumor Samples |
| --- | --- | --- | --- |
| Liver-HCC | LIVER | Liver hepatocellular carcinoma | 306 |
| Panc-AdenoCA | PANCREAS | Pancreatic adenocarcinoma | 235 |
| Breast-AdenoCA | BREAST | Breast adenocarcinoma | 198 |
| Prost-AdenoCA | PROSTATE GLAND | Prostate adenocarcinoma | 189 |
| CNS-Medullo | BRAIN, & CRANIAL NERVES, & SPINAL CORD, | Medulloblastoma | 146 |
| Kidney-RCC | KIDNEY | Renal cell carcinoma (proximal tubules) | 143 |
| Ovary-AdenoCA | OVARY | Ovarian adenocarcinoma | 112 |
| Skin-Melanoma | SKIN | Skin melanoma | 106 |
| Lymph-BNHL | LYMPH NODES | Mature B-cell lymphoma | 105 |
| Eso-AdenoCA | ESOPHAGUS | Esophageal adenocarcinoma | 98 |
| Lymph-CLL | BLOOD, BONE MARROW, & HEMATOPOIETIC SYS | Chronic lymphocytic leukemia | 95 |
| CNS-PiloAstro | BRAIN, & CRANIAL NERVES, & SPINAL CORD, | Pilocytic astrocytoma | 89 |
| Panc-Endocrine | PANCREAS | Pancreatic neuroendocrine tumor | 85 |
| Stomach-AdenoCA | STOMACH | Gastric adenocarcinoma | 70 |
| Head-SCC | GUM, FLOOR OF MOUTH, & OTHER MOUTH | Head/neck squamous cell carcinoma | 57 |
| ColoRect-AdenoCA | LARGE INTESTINE, (EXCL. APPENDIX) | Colorectal adenocarcinoma | 52 |
| Lung-SCC | LUNG & BRONCHUS | Lung squamous cell carcinoma | 48 |
| Thy-AdenoCA | THYROID GLAND | Thyroid adenocarcinoma | 48 |
| Myeloid-MPN | BLOOD, BONE MARROW, & HEMATOPOIETIC SYS | Myeloproliferative neoplasm | 46 |
| Kidney-ChRCC | KIDNEY | Renal cell carcinoma (distal tubules) | 45 |
| Bone-Osteosarc | BONES & JOINTS | Sarcoma, bone | 44 |
| CNS-GBM | BRAIN, & CRANIAL NERVES, & SPINAL CORD, | Diffuse glioma | 41 |
| Uterus-AdenoCA | UTERUS, NOS | Uterine adenocarcinoma | 40 |
| Lung-AdenoCA | LUNG & BRONCHUS | Lung adenocarcinoma | 38 |
|  |  |  | **2436** |

### Classification using Single Mutation Feature Types

To determine the predictive value of different mutation features, we trained and evaluated a series of tumour type classifiers based on single categories of feature derived from the tumour mutation profile. For each feature category we developed a random forest (RF) classifier (Online Methods). Each classifier’s input was the mutational feature profile for an individual tumour specimen, and its output was the probability estimate that the specimen belongs to the type under consideration. Each classifier was trained using a randomly selected set of 75% of samples drawn from the corresponding tumour type. To determine the most likely type for a particular tumour sample, we applied its mutational profile to each of the 24 type-specific classifiers, and selected the type whose classifier emitted the highest probability. To evaluate the performance of the system, we applied stratified four-fold cross-validation by training on three quarters of the data set and testing against each of the other quarter specimens. We report overall accuracy as well as recall, precision and the F1 score using the average of all four test data sets (see Online Methods for cross-validation methodology and definitions of terms).

We selected a total of seven mutational feature types spanning three major categories (Table 2):

**Table 2:** WGS feature types used in classifiers

| Feature Category | Feature Type | Feature Count | Description |
| --- | --- | --- | --- |
| **Mutation Distribution** | SNV-BIN | 2897 | Number of SNVs per 1 Mbp bin, and per chromosome,, normalized against total number of SNVs per sample |
|  | CNA-BIN | 2826 | Number of CNAs per 1 Mbp bin |
|  | SV-BIN | 2929 | Number of SVs per 1 Mbp bin, and per chromosome,, normalized against total number of SV per sample |
|  | INDEL-BIN | 2757 | Number of SNVs per 1 Mbp bin, and per chromosome,, normalized against total number of INDEL per sample |
| **Mutation Type** | MUT-WGS | 150 | Type of single nucleotide substitution,, double, and triple nucleotide substitution (plus its adjacent nucleotide neighbors) |
| **Driver Gene/Pathway** | GEN | 554 | Presence of an impactful mutation in a suspected driver gene |
|  | MOD | 1865 | Presence of an impactful mutation in a gene belonging to a suspected driver pathway |

- *Mutation Distribution.* The somatic mutation rate in cancers varies considerably from one region of the genome to the next.^2^ In whole genome sequencing, a major covariate of this regional variation in whole genome sequences is the epigenetic state of the tumour’s cell of origin, with 74-86% of the variance in the mutation density being explained by histone marks and other chromatin features related to open versus closed chromatin^12^. This suggests that tumours sharing similar cells of origin will have a similar topological distribution of mutations across the genome. To capture this, we divided the genome into ∼3000 1 Mbp bins across the autosomes (excluding sex chromosomes) and created features corresponding to the number of somatic mutations per bin normalized to the total number of somatic mutations. Mutation rate profiles were created independently for somatic substitutions (SNV), indels, somatic copy number alterations (CNA), and other structural variations (SV). Note that the vast majority of variants, e.g., at least 99% of the SNVs in nearly all samples, used for this analysis are non-functional passenger mutations. See Campbell *et al*^4^ and Wala *et al.*^14^ for descriptions of point and structural variations in the PCAWG dataset.
- *Mutation Type.* The type of the mutation and its nucleotide neighbors, for example G{C>T}C, is an indicator of the exposure history of the cell of origin to extrinsic and endogenous factors that promote mutational processes^15^. This in turn can provide information on the etiology of the tumour. For example, skin cancers have mutation types strongly correlated with UV light-induced DNA damage. Reasoning that similar tumour types will have similar mutational exposure profiles, we generated a series of features that represented the normalized frequencies of each potential nucleotide change in the context of its 5’ and 3’ neighbors. Like the mutation distribution, the variants that contribute to this feature category are mostly passengers. Readers are referred to Alexandrov *et al*^16^ for more information on signature analysis in the PCAWG data set.
- *Driver Gene/Pathway.* Some tumour types are distinguished by high frequencies of alterations in particular driver genes and pathways. For example, melanomas have a high frequency of BRAF gene mutations^17^, while pancreatic cancers are distinguished by KRAS mutations^8^. We captured this in two ways: (1) whether a gene is affected by a driver event as determined by the PCAWG Cancer Drivers Working Group^18^, and (2) whether there was an impactful coding mutation in any gene belonging to a known or suspected driver pathway (also see Reyna *et al*^19^ for cancer pathway analysis performed by the PCAWG Pathway and Networks Working Group). We counted driver events affecting protein-coding genes, long noncoding RNAs and micro-RNAs, but did not attempt to account for alterations in cis-regulatory regions. In all we created ∼2000 driver pathway-related features describing potential gene and pathway alterations for each tumour.

The accuracy of individual RF classifiers ranged widely across tumour and feature categories, with a median F1 (harmonic mean of recall and precision) of 0.42 and a range from 0.00 to 0.94 (Figure 1a,b, Supplementary Figure 1, Supplementary Tables 2). Nine tumour types had at least one well-performing classifier that achieved an F1 of 0.80: CNS-GBM, CNS-PiloAstro, Liver-HCC, Lymph-BNHL, Kidney-RCC, Myeloid-MPN, Panc-AdenoCA, Prost-AdenoCA, Skin-melanoma. Five classifiers performed poorly, with no classifier achieving an accuracy greater than 0.6: Bone-Osteosarc, Head-SCC, Stomach-AdenoCA, Thy-AdenoCA and Uterus-AdenoCA. The remaining eight tumour types had classifiers achieving F1s between 0.60 and 0.80.

**Figure 1.**
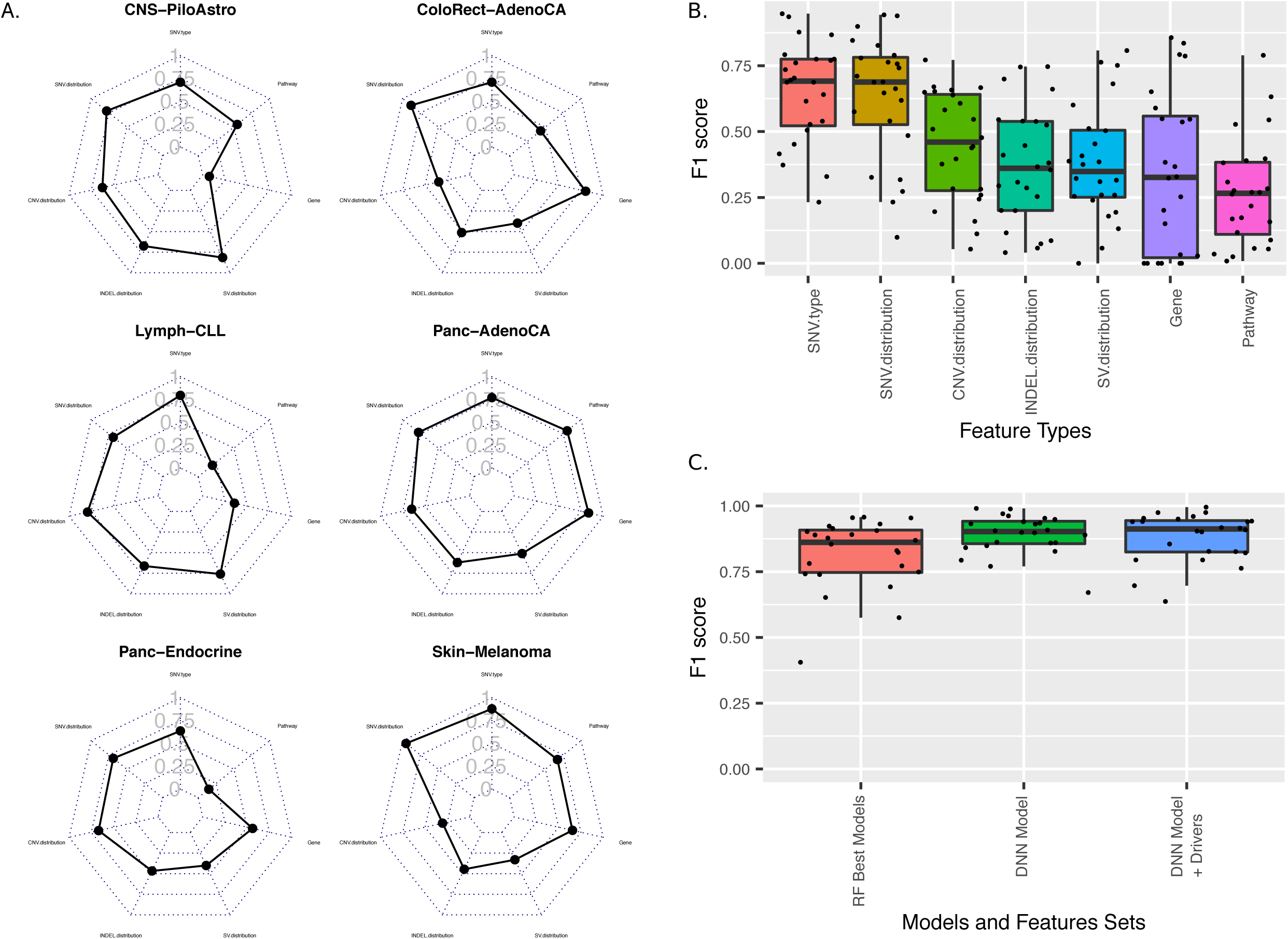
Comparison of tumour type classifiers using single and multiple feature types. (A) Radar plots describing the cross-validation derived accuracy (F1) score of Random Forest classifiers trained on each of 7 individual feature categories, across six representative tumour types. (B) Summary of Random Forest classifier accuracy (F1) trained on individual feature categories across all 24 tumour types. (C) Accuracy of classifiers trained on multiple feature categories. *RF Best Models* corresponds to the cross-validation F1 scores of Random Forest classifiers trained on the three best single feature categories for each tumour type. *DNN Model* shows the distribution of F1 scores for held out samples for a multi-class neural network trained using passenger mutation distribution and type. *DNN Model + Drivers* shows F1 scores for the neural net when driver genes and pathways are added to the training features.

The highest accuracies were observed for features related to mutation type and distribution (Figure 1b). Contrary to our expectations, altered driver genes and pathways were poor discriminatory features. Whereas both SNV type and distribution achieved median F1 scores of ∼0.7, RF models built on driver gene or pathway features achieved median F1s of 0.33 and 0.27 respectively. Only Panc-AdenoCA, Kidney-RCC, Lymph-BNHL and ColoRect-AdenoCA exceeded F1s greater than 0.75 on RF models built from gene or pathway-related features, but we note that even in these cases, the mutation type and/or distribution features performed equally well.

### Classification using Combinations of Mutation Feature Types

We next asked whether we could improve classifier accuracy by combining features from two or more categories. We tested both Random Forest (RF) and multi-class Deep Learning/Neural Network (DNN)-based models (Online Methods), and found that overall the DNN-based models were more accurate than RF models across a range of feature category combinations (median F1=0.86 for RF, F1=0.90 for DNN, p<1.2e-7 Wilcoxon Rank Sum Test; Figure 1C). For the DNN-based models, overall accuracy was the highest when just the topological distribution and mutation type of SNVs were taken into account. Adding gene and/or pathway features slightly reduced classification accuracy; using only gene and pathway features greatly reduced classifier performance. We did not investigate the effect of training the DNN on CNV or SV features as these mutation types were not uniformly available in the validation data sets (see below).

Figure 2 shows a heatmap of the DNN classifier accuracy when tested against held out tumours (mean of 10 independently-built models). Overall, the accuracy for the complete set of 24 tumour types was 91% (classification accuracy), but there was considerable variation for individual tumours types (Supplementary Table 3). Recall (also known as sensitivity) ranged from 0.61 (Stomach-AdenoCA) to 0.99 (Kidney-RCC). Precision (similar to specificity but is sensitive to the number of positives in the data set) was comparable, with rates ranging from (Stomach-AdenoCA) to 1.00 (CNS-GBM, Skin-Melanoma, and Liver-HCC). Twenty-one of 24 tumour types achieved F1s greater than 0.80, including 8 of the 9 of the types that met this threshold for RF models built on single feature categories. The three worst-performing tumour types were CNS-PiloAstro (mean F1 0.79 across 10 independently-trained DNN models), Lung-AdenoCA (F1 0.77) and Stomach-AdenoCA (F1 0.67).

**Figure 2.**
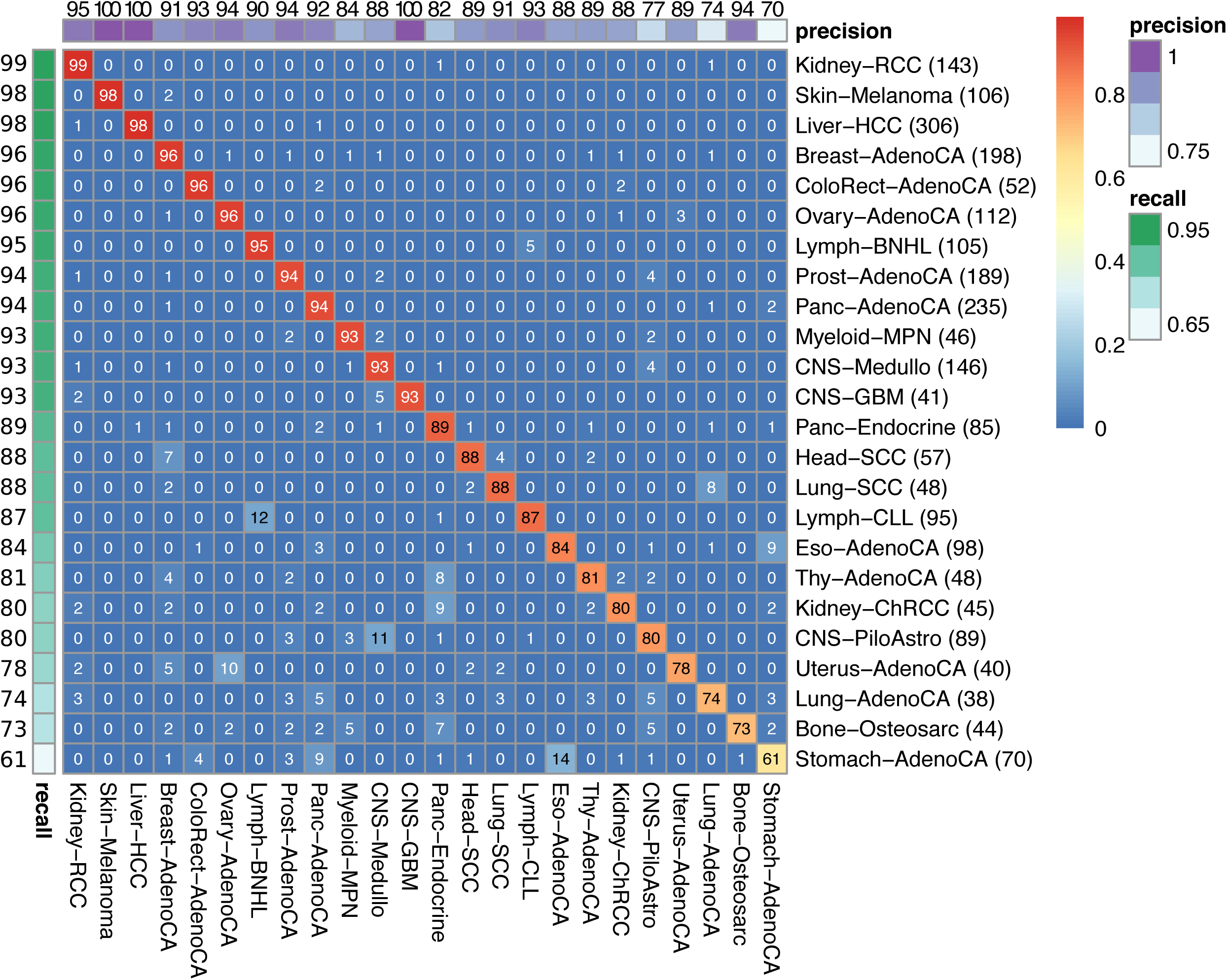
Heatmap displaying the accuracy of the merged classifier using a held out portion of the PCAWG data set for evaluation. Each row corresponds to the true tumor type; Columns correspond to the class predictions emitted by the DNN. Cells are labeled with the percentage of tumors of a particular type that were classified by the DNN as a particular type. The recall and precision of each classifier is shown in the color bars at the top and left sides of the matrix. All values represent the mean of 10 runs using selected data set partitions.

We investigated the effect of the training set size on classifier accuracy (Figure 3a). Tumour types with fewer than 100 samples in the data set were more likely to make incorrect predictions, and tumour types with large numbers of samples were among the top performers. However, several tumour types including ColoRect-AdenoCA (N=52), Lung-SCC (N=48) and CNS-GBM (N=41) achieved excellent predictive accuracy despite having small training sets.

**Figure 3.**
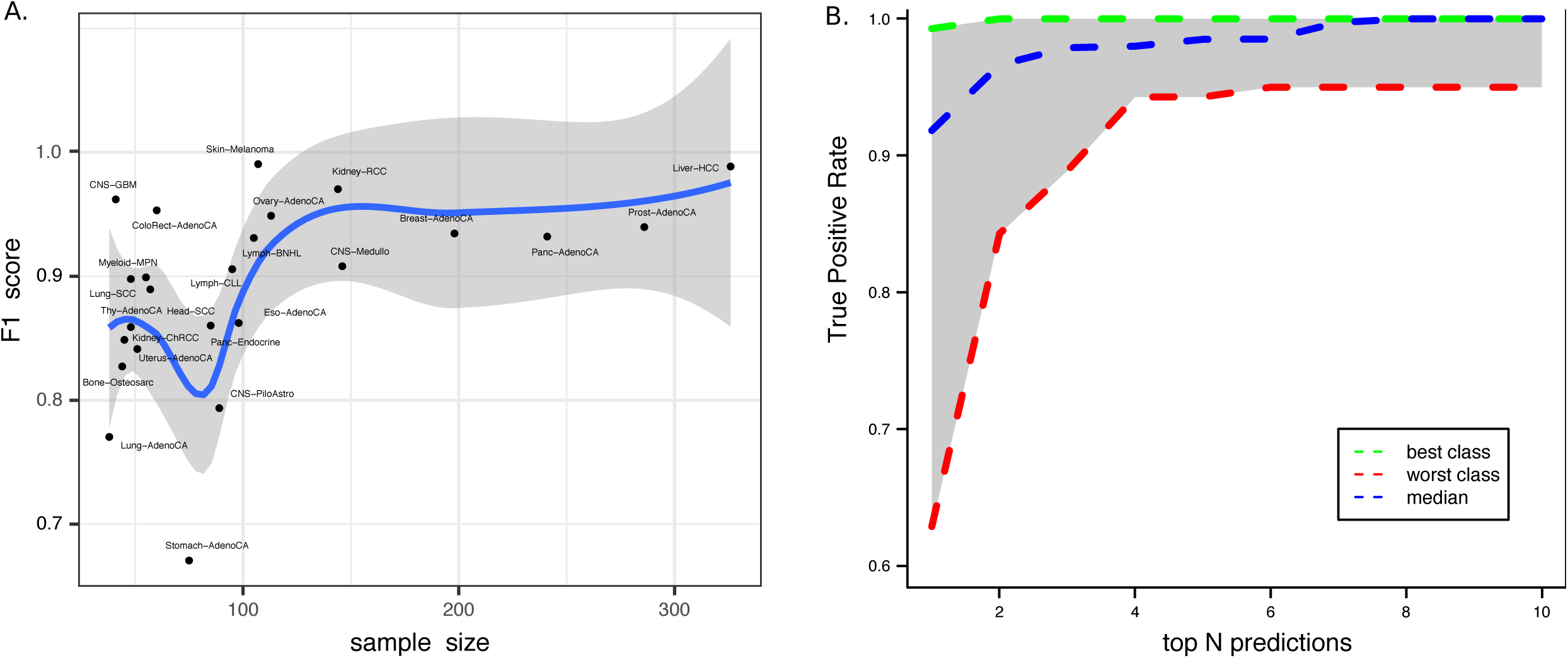
(A) Relationship between training set size and prediction accuracy of the DNN. The blue line represents a regression line fit using LOESS regression. The grey area represents a 95% confidence interval for the regression function. (B) Frequency with which the correct tumour type was contained within the DNN’s top ranked predictions.

The DNN emits a softmax output that can be interpreted as the probability distribution of the tumour sample across the 24 cancer types. We ordinarily select the highest probability tumour type as the classifier’s choice. If instead we asked how often the correct type is contained among the top N ranked probabilities, we find that the worst performing tumour type (Stomach-AdenoCA) achieved an true positive rate of of 0.88 for placing the correct tumour type among the top ranked three choices, and that the average true positive rate across all tumour types for this task was 0.98 (Figure 3b).

### Patterns of Misclassification

Misclassifications produced by the DNN in many cases seem to reflect shared biological characteristics of the tumours. For example, the most frequent classification errors for Stomach-AdenoCA samples were to two other upper gastrointestinal tumours, esophageal adenocarcinoma (Eso-AdenoCA, 14% misclassification rate), and pancreatic ductal adenocarcinoma (Panc-AdenoCA, 9%). These three organs share a common developmental origin in the embryonic foregut and may share similar epigenetic profiles. We also speculate that the high rate of confusion between gastric and esophageal cancers might be due to similar mutational exposures among the two sites: a subset of C->A, C->G substitutions are commonly seen in stomach and esophageal (but not pancreatic) cancers and comprise Signature 17 in the COSMIC catalogue of mutational signatures^20^. To test this, we assessed the effect of training the DNN with mutation distribution alone, excluding mutation type features (Supplementary Figure 2). Using just passenger mutation distribution, the overall F1 for stomach tumours increased by 4%, supporting the idea that part of the error is due to shared mutational signatures among stomach and esophageal cancer. Another possible explanation for the frequent misclassification of gastric and esophageal tumours is that some of the tumours labeled gastric arose at the gastroesophageal junction (GEJ), which some consider to be a distinct subset of esophageal tumours^21^.

Other common misclassification errors include misclassification of 12% of chronic lymphocytic leukemia (Lymph-CLL) samples as B-cell non-hodgkin’s lymphoma (Lymph-BNHL). Both tumours are derived from the B-cell lymphocyte lineage, and likely share a similar cell of origin. Another pattern was occasional misclassifications among the three types of brain tumour CNS-GBM, CNS-Medullo, and CNS-PiloAstro, all three of which are derived from various glial lineages. We speculate that these errors are again due to similarities among the cells of origin of these tissues.

Of note is that the DNN was able to accurately distinguish among several tumour types that arise from the same organ. Renal cell carcinoma (Kidney-RCC) and chromophobe renal carcinoma (Kidney-ChRCC), were readily distinguished from each other, as were the squamous and adenocarcinoma forms of non-small cell lung cancer (Lung-SCC, Lung-AdenoCA), and the exocrine and endocrine forms of pancreatic cancer (Panc-AdenoCA, Panc-Endocrine). The misclassification rate between Lung-SCC and Lung-AdenoCA was just 8%, and all other pairs had misclassification rates of 2% or lower. This is in keeping with a model in which major histological subtypes of tumours often reflect different cells of origin.

### Validation on an Independent Collection of Primary Cancer Whole Genomes

A distinguishing characteristic of the PCAWG data set is its use of a uniform computational pipeline for sequence alignment, quality filtering, and variant calling. In real world settings, however, the data set used to train the classifier may be called using a different set of algorithms than the test data. To assess the accuracy of DNA-based tumour identification when applied in this setting, we applied the classifier trained on PCAWG samples to an independent validation set of 1,436 cancer whole genomes assembled from a series of published non-PCAWG projects. The validation set spans 14 distinct tumour types assembled from 21 publications or databases (Supplementary Table 4). We were unable to collect sufficient numbers of independent tumour genomes representing nine of the 24 types in the merged classifier, including colorectal cancer, thyroid adenocarcinoma and lung squamous cell carcinoma. SNV coordinates were lifted from GRCh38 to GRCh37 when necessary, but we did not otherwise process the mutation call sets. With the exception of a set of liver cancer (Liver-HCC) samples in the validation set, which is discussed below, a comparison of the mutation load among each tumour type cohort revealed no significant differences between the PCAWG and validation data sets (Supplementary Figure 3).

The DNN classifier recall for the individual tumour types included in the validation data set ranged from 0.41 to 0.98, and the precision ranged from 0.36 to 1.0 (Figure 4a), achieving an overall accuracy of 82% for classification across the multiple types. In general, the tumour types that performed the best within the PCAWG data set were also the most accurate within the validation, with Panc-AdenoCA, Skin-Melanoma, Kidney-RCC, Ovary-AdenoCA and Breast-AdenoCA tumour types all achieving greater than 80% accuracy. The Lymph-CLL, Liver-HCC, Eso-AdenoCA, CNS-GBM and CNS-Medullo were poorly predicted with recalls below 70%, and the remaining types had intermediate accuracies.

**Figure 4.**
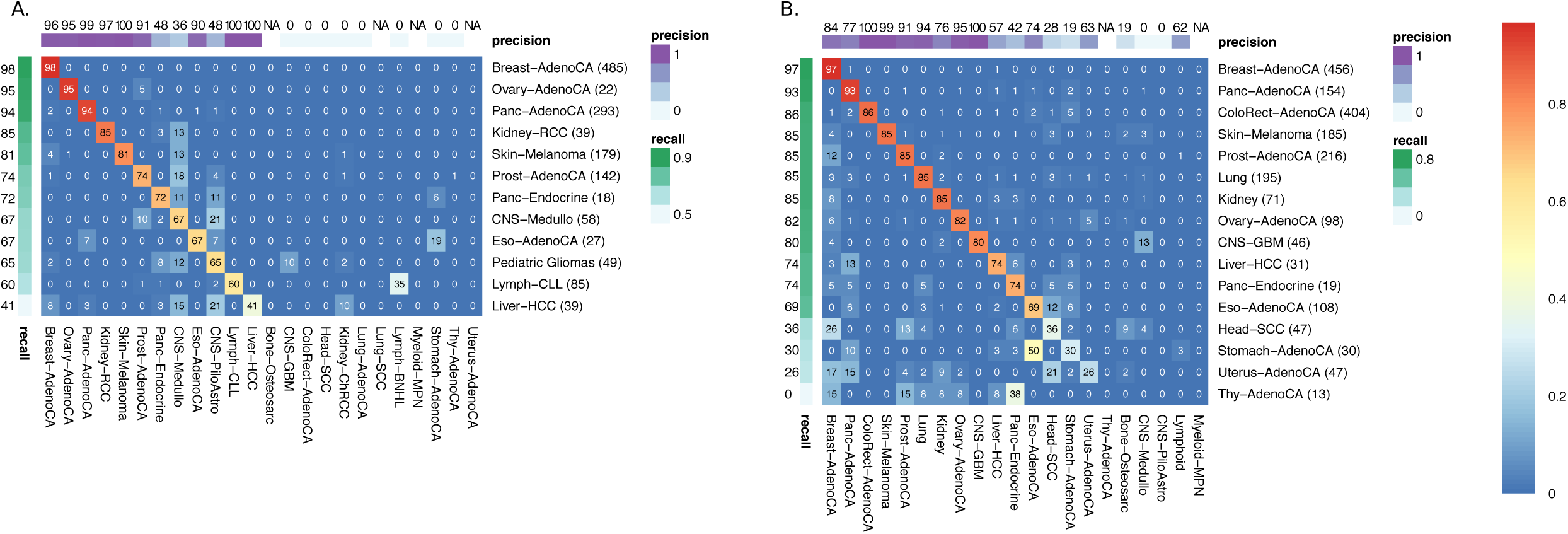
Prediction accuracy for the DNN against two independent validation data sets. (A) Primary tumours. (B) Metastatic tumours. Details on the validation data sets are described in Results. The format of the heatmap is the same as described in Figure 2.

The majority of classification errors observed in the primary tumour validation set mirrored the patterns of misclassifications previously observed within the PCAWG samples, with the exception that Liver-HCC cases were frequently misclassified as CNS-PiloAstro (21%) and CNS-Medullo (15%). We believe this case to be due to a lower than expected mutation burden in the liver tumours from the validation set (median 3202 SNVs per sample in validation set vs 22230 SNVs per sample in the PCAWG training set; P<1.5e-15 by Wilcoxon Rank Sum Test; Supplementary Figure 3). This mutation load is more similar to the rates observed in CNS-PiloAstro (median 344 per sample) and CNS-Medullo (median 2330 per sample) among the PCAWG samples, and might suggest poor coverage of Liver-HCC or another sequencing/analysis artifact in the validation set.

We were initially puzzled that a set of 49 validation data set samples that were identified as “CNS glioma” overwhelmingly matched to the pediatric piloastrocytoma model rather than to the CNS-GBM model. However, on further investigation, we discovered that these samples represent a mixture of low- and high-grade pediatric gliomas, including piloastrocytomas^22-24^. The SNV mutation burden of these pediatric gliomas is also similar to CNS-PiloAstro and significantly lower than adult CNS-GBM (Supplementary Figure 3).

### Validation on Metastatic Tumors

To evaluate the ability of the classifier to correctly identify the type of the primary tumour from a metastatic tumor sample, we developed an independent validation data set that combined a published series of 92 metastatic Panc-AdenoCA^25^ with an unpublished set of 2,028 metastatic tumours from known primaries across 16 tumour types recently sequenced by the Hartwig Medical Foundation (HMF)^26^, resulting in a combined set of 2,120 samples across 16 tumour types (Supplementary Table 4). All metastatic samples were subjected to paired-end WGS sequencing of tumour and normal at a tumour coverage of at least 65x, but the computational pipelines used for alignment, quality filtering, and SNV calling were different from those used for PCAWG. The rules for matching classifier output to the validation set class labels were developed in advance of the experiment, and the DNN classifier was applied to the molecular data from the validation set in a blind fashion.

When the DNN classifier was applied to these metastatic samples it achieved an overall accuracy of 85% for identifying the type of the known primary (Figure 4b), which is similar to its performance on the validation primaries. Nine of the tumour types in the metastatic set achieved recall rates of 0.80 or higher, including Breast-AdenoCA (0.97), Panc-AdenoCA (0.93), and ColoRect-AdenoCA (0.86). On the other end of the spectrum, four tumour types failed to achieve a recall of at least 0.50: Head-SCC (0.36), Stomach-AdenoCA (0.30), Uterus-AdenoCA (0.26) and Thyroid-AdenoCA (0.0). Overall, the patterns of misclassification were similar to what was seen within PCAWG. For example, the gastric cancers were misclassified as esophageal tumours 50% of the time.

In contrast to the other tumour types, metastatic thyroid adenocarcinoma was a clear outlier. In this case, the DNN was unable to correctly identify any of the 13 metastatic samples, classifying them instead as other tumour types such as Panc-Endocrine, Prost-AdenoCA or Breast-AdenoCA. We lack information on the histological subtype of the metastatic thyroid tumours in the HMF data set, but speculate that the metastatic thyroid tumours in this set are enriched in more aggressive histological subtypes than the PCAWG primaries, which are exclusively of low-grade papillary (N=31), papillary-follicular (N=18) and papillary-columnar (N=1) types.

The HMF data set also included 62 CUPs tumours. While we do not know the corresponding primary for these samples, we did attempt to classify them (Supplementary Table 5). The CUPs cases were most frequently classified as Panc-AdenoCA (N=9; 15%), Lung-AdenoCA (N=9; 15%) and Liver-HCC (N=8; 13%). Reassuringly, despite the fact that information on the sex chromosomes were **not** used by the classifier, all the CUPs tumours classified as gynecological tumours (Breast-AdenoCA, N=5; Uterus-AdenoCA, N=2) came from female patients. Interestingly, while the classifier made a low probability prediction for one female patient as Prost-AdenoCA, the second-best prediction was for Uterus-AdenoCA, and its probability was almost identical to the higher ranked prediction (0.26 vs 0.27).

### Code Availability

The code developed for training and testing the classifier, along with documentation and trained models for the 24 tumour types are available from GitHub at https://github.com/ICGC-TCGA-PanCancer/TumorType-WGS.git. The code is distributed under the Apache Version 2.0 Open Source license (https://www.apache.org/licenses/LICENSE-2.0).

## Discussion

Cancer of unknown primary site (CUPS) is a heterogeneous set of cancers diagnosed when a patient presents with metastatic disease, but despite extensive imaging, pathological and molecular studies the primary cannot be determined.^11^ CUPS accounts for 3-5% of cancers, making it the seventh to eighth most frequent type of cancer and the fourth most common cause of cancer death^27^. Even at autopsy, the primary cannot be identified roughly 70% of the time^28^, suggesting regression of the primary in many CUPS cases. CUPS is a clinical dilemma, because therapeutic options are largely driven by tissue of origin, and site-directed therapy is more effective than broad-spectrum chemotherapy^29^. A related diagnostic challenge arises, paradoxically, from the medical community’s success in treating cancers and the rising incidence of second primary cancers, now estimated at roughly 16% of incident cancers^30^. Pathologists are often asked to distinguish a late metastatic recurrence of a previously treated primary from a new unrelated primary. However, histopathology alone may be inaccurate at identifying the site of origin of metastases. In one study^31^, pathologists who were blinded to the patient’s clinical history were able to identify the primary site of a metastasis no more than 49% of the time when given a choice among 11 adenocarcinomas. When asked to rank their guesses, the correct diagnosis was among the top 3 choices just 76% of the time.

In this paper, we used the largest collection of uniformly processed primary cancer whole genomes assembled to date to develop a supervised machine learning system capable of accurately distinguishing 24 major tumour types based solely on features that can be derived from DNA sequencing. The accuracy of the system overall was 91%, with 20 of the 24 tumour types achieving an F1 score of 0.83 or higher. When the tumour type predictions were ranked according to their probability scores, the correct prediction was found among the top three rankings 98% of the time.

To independently validate the classifier, we assessed it using a set of 1,436 primary tumours that had been subjected to WGS by independent groups. This validation set represented 12 of the PCAWG tumour types and achieved an overall predictive accuracy of 85%. Further validation using an independent set of 2120 metastatic tumours corresponding to 16 known primary sites achieved an overall accuracy of 82%. Some of the reduction we observed in the classifier’s prediction accuracy when applied to the independent data sets was likely due to their differing somatic mutation-calling pipelines, which use different quality-control filters, genome builds and SNV callers from those applied to PCAWG samples. In support of this conclusion, we found that the some of the worst performing tumour types were those in which the mutation load in the validation sets deviated widely from the load in the corresponding PCAWG tumour type.

The regional distribution of somatic passenger mutations across the genome was the single most predictive class of feature, followed by the distribution of mutation types. The regional density of somatic mutations is thought to reflect chromatin accessibility to DNA repair complexes, which in turn relates to the epigenetic state of the cancer’s cell of origin. The DNN’s predictive accuracy is therefore largely driven by a cell of origin signal, aided to a lesser extent by signatures of exposure. The observation that the classifier was able to identify the site of origin for a series of metastatic tumours with the same or better accuracy as its performance on primaries suggests that the cell of origin and exposure signals are already established in the early cancer (or its precursor cell) and are not masked by subsequent mutations that occur during tumour evolution.

Unexpectedly, the distribution of functional mutations across driver genes and pathways were poor predictors of tumour type in all but a few tumour types (e.g. pancreatic adenocarcinoma). This surprising finding may be explained by the observation that there are relatively few driver events per tumour (mean 4.6 events per tumour^4^), and affect a set of common biological pathways related to the “hallmarks of cancer.”^32^ This finding may also explain the observation that automated prediction of tumor type by exome or gene panel sequencing has so far met with mixed success (see below).

There was considerable variability in the classification accuracy among tumour types. In most cases tumour types that were frequently confused with each other had biological similarities such as related tissues or cells of origin. Technical issues that could degrade predictive accuracy include uneven sequencing coverage, low sample purity, inadequate numbers of samples in the training set, and tumour type heterogeneity. A larger collection of tumours with WGS would allow us to improve the classifier accuracy as well as to train the classifier to recognize clinically-significant subtypes of tumours, such as the basal form of breast cancer.

There are other ways of identifying the site of origin of a tumor. In cases in which the tumour type is uncertain pathologists frequently apply a series of antibodies to tissue sections to detect tissue-specific antigens via immunohistochemistry (IHC). The drawback of IHC is that it requires manual interpretation, and the decision tree varies according to the differential diagnosis^33^. Furthermore, IHC is known to be confounded by the loss of antigens in poorly differentiated tumours^34^. In principle, tumour differentiation state should not impact the performance of our classifier because it relies on the distribution of passenger mutations, most of which are already established at the time of tumour initiation. Because of the many different grading systems applied across the PCAWG set a direct test of this notion is difficult, but we are reassured that the independent set of metastases, which frequently represent a higher grade than the primary, performed as well as the external primary tumour validation set.

An alternative to IHC is molecular profiling of tumors using mRNA or miRNA expression, and several commercial systems are now available to identify the tissue of origin using microarray or qRT-PCR assays^35-37^. A recent comparative review^35^ of five commercial expression-based kits reported overall accuracies between 76 and 89%; the number of tumor types recognized by each system ranges from six to 47 with accuracy tending to decrease as the number of discriminated types increases.

Patterns of DNA methylation are also strongly correlated with the tissue of origin. A recent report^38^ demonstrated highly accurate classification of more than 70 central nervous system tumour types using a Random Forest classifier trained on methylation array data. Another recent report^39^ showed that an immunoprecipitation-based protocol can recover circulating tumour DNA from patient plasma and accurately distinguish among three tumour types (lung, pancreatic and AML) based on methylation patterns.

Previous work in the area of DNA-based tumour type identification has used targeted gene panel^40^ and whole exome^41-43^ sequencing strategies. The targeted gene-based approach described in Tothill^40^ is able to discriminate a handful of tumour types that have distinctive driver gene profiles, and can identify known therapeutic response biomarkers, but does not have broader applicability to the problem of tumour typing. In contrast, the whole exome sequencing approaches reported by Chen^41^, Soh^42^ and Marquard^43^ each used machine learning approaches to discriminate among 17, 28, and 10 primary sites, respectively, achieving overall accuracies of 62%, 78% and 69%. Interestingly, all three papers demonstrated that classifiers built on multiple feature categories outperformed those built on a single type of feature, consistent with our findings. We demonstrate here that the addition of whole genome sequencing data substantially improves discriminative ability over exome-based features. It is also worth noting that Soh^41^ was able to achieve good accuracy using SNVs and CNAs spanning just 50 genes, suggesting that it may be possible to retain high classifier accuracy while using mutation ascertainment across a well-chosen set of whole genomic regions.

In practical terms, whole genome sequencing and analysis of a cancer tumor/normal pair costs $3000-4000 USD and can be completed in 2-3 weeks (P. Krzyzanowski, OICR Genomics Program, and E. Cuppen, Hartwig Medical Foundation, personal communications). As the price continues to drop, there is an accelerating trend to apply genome sequencing to routine cancer care in order to identify actionable mutations and to test for the presence of predictive biomarkers, and some centres are now able to turn around NGS-based sequencing results in 24 hours (David Louis, personal communication). An example of the trend is the National Health Service of the UK, which recently announced a plan to apply WGS routinely to cancer patients^44^. Given the increasing likelihood that many or most cancers will eventually have genomic profiling, it is attractive to consider the possibility of simultaneously deriving the cancer type using an automated computational protocol. This would serve as an adjunct to histopathological diagnosis, and could also be used as a quality control check to flag the occasional misdiagnosis or to find genetically unusual tumours. More forward-looking is the prospect of accurately determining the site of origin of circulating cell-free tumour DNA detected in the plasma using so-called “liquid biopsies”^45^, possibly in conjunction with methylome analysis.^38,39^ As genome sequencing technologies continue to increase in sensitivity and decrease in cost, there are realistic prospects for blood tests to detect early cancers in high risk individuals^46^. The ability to suggest the site and histological type of tumours detected in this way would be invaluable for informing the subsequent diagnostic workup.

In summary, this is the first study to demonstrate the potential of whole genome sequencing to distinguish major cancer types on the basis of somatic mutation patterns alone. Future studies will focus on improving the classifier performance by training with larger numbers of samples, subdividing tumour types into major molecular subtypes, adding new feature types, and adapting the technique to work with clinical specimens such as those from formalin-fixed, paraffin-embedded biopsies and cytologies.

## Acknowledgements

We would like to thank Irina Kalatskaya, Quang Trinh, Jared Simpson, Katie Hoadley and David Louis for their helpful comments during preparation of this manuscript. We also gratefully acknowledge the assistance of Drs. Ludmil B. Alexandrov, Mi Ni Huang, Arnoud Boot, Steven Gallinger, Julie Wilson, Haiko J. Bloemendal, Laurens Beerepoot, Steven G. Rozen and Michael R. Stratton in providing independent WGS primary and metastatic tumour SNV profiles used for validation. We also thank WJ, LS, and QM are supported by funding from the Province of Ontario, Canada. QM’s research was supported by a gift from NVIDIA foundation, an advised fund of the Silicon Valley Community Foundation.

