## Supplementary material for "A deep learning system can accurately classify primary and metastatic cancers based on patterns of passenger mutations": Online Methods

*PCAWG Training and Testing Data Set*

All variant call data were downloaded from Synapse (<https://www.synapse.org/#!Synapse:syn2351328/wiki/62351>; the “syn” numbers that follow refer to Synapse data set IDs. Consensus Somatic SNV and INDEL (syn7118450) file covers 2778 whitelisted samples from 2583 donors. Consensus SV calls from the PCAWG Structural Variation Working Group were downloaded in VCF format (syn7596712). Ploidy and purity information are from the PCAWG Evolution and Heterogeneity Working Group (syn8042880, syn8272483) and driver events were called by the PCAWG Drivers and Functional Interpretation Group (syn9757986). Tumour histological classifications were reviewed and assigned by the PCAWG Pathology and Clinical Correlates Working Group (annotation version 6, August 2016; syn7253568). For model training, we first removed all samples that had been flagged as exhibiting microsatellite instability (MSI) by the PCAWG Technical Working Group. In a small number of cases, the same donor contributed both primary and metastatic tumour specimens to the PCAWG data set. In these cases we used only the primary tumor for training and evaluation, except for the case of the small cohort of myeloproliferative neoplasms (Myeloid-MPN; N=55 samples), for which multiple primary samples were available. In this case, we used up to two samples per donor and partitioned the training and testing sets to avoid having the same donor appear more than once in any training/testing set trial (see Supplementary Table 1 for the complete list of tumour specimens).

*Feature Sets*

Mutational type features are based on all point substitutions (single nucleotide variations; SNVs). For each sample, SNVs are categorized across the six possible single nucleotide changes (A->C, A->G, A->T, C->A, C->G, C-T), the 48 possible nucleotide changes plus their 5' or 3' flanking base, and the 96 possible nucleotide changes plus both flanking nucleotides. This generates 150 mutational type features in total. The counts in each category are then normalized to the total number of SNVs in the sample.

Mutational distribution features are the number of SNVs, small indels, structural variation (SV) breakpoints, and somatic copy number variations (CNVs) in each 1 megabase bins across the genome. The total number of SNV, indel and SV counts in each bin were normalized to the total number of the corresponding mutational events across the genome. In addition, we generated the following features: (1) the total numbers of each type of mutational event per genome; (2) the number of each type of mutational event per chromosome, normalized by chromosome length; (3) sample purity values; and (4) sample ploidy. In total, there are 2897 SNV+indel, 2826 CNV, and 2929 SV features. For the initial selection of feature types, we tested all mutational distribution features. However, the final neural network used SNV features only.

Driver gene and pathway features were derived from the driver event list generated by the PCAWG Drivers and Functional Interpretation Working Group (https://www.biorxiv.org/content/early/2017/12/23/237313). This list contains driver events in coding genes, as well as events that affect miRNA and lncRNAs. We generated a boolean matrix from the list in which each row is a tumour sample and each column is a driver event. To mutations to pathways, we selected any non-synonymous SNV affecting a gene in a pathway, regardless of its putative driver status. These SNVs were then assigned to 1,865 pathways from the Reactome resource (<http://www.reactome.org>, version 58)^1^. A pathway feature was scored as positive if it contained at least one driver gene. Because a gene may be contained within more than one pathway, it is possible for a single driver gene event to generate two or more positive pathway features.

*Independent validation data set: Primary and Metastatic Tumours*

To independently validate the neural network-based classifier, we assembled several sets of tumours that had been subject to whole genome sequencing outside of PCAWG (Supplementary Table 4).

The primary tumour validation data set consisted of 1236 primary tumours contributed by colleagues participating in the PCAWG Mutational Signatures Working Group and described in^2^. These represent 12 tumour types overlapping with PCAWG types collected from a variety of published studies, non-PCAWG donors submitted to the ICGC data portal (<http://dcc.icrg.org>), and donors present in the COSMIC database (<http://cancer.sanger.ac.uk/cosmic>). These independent primaries were supplemented using WGS data from 200 advanced primary pancreatic ductal adenocarcinomas (Panc-AdenoCA) derived from the COMPASS Trial^3^ and used with the gracious permission of Dr. Steven Gallinger. In all, the primary tumour validation set contained 1436 primary tumour samples across 12 tumour types. Only tumour types with 10 or more representatives were used for testing.

The metastatic tumour validation data set was derived from SNV calls on 2028 metastatic tumours across 16 tumour types. They are a subset of 2090 total samples provided by Dr. Edwin Cuppen with matched PCAWG histology subtypes and are described in Supplementary Table 4 and Priestley*, et al.*^4^ We supplemented this set with 92 metastatic pancreatic ductal adenocarcinomas to the liver from the COMPASS Trial, for a total of 2120 metastatic tumours. As for the primaries, only tumour types with 10 or more representatives were tested.

Although the sequencing technologies and genome coverage are comparable among the PCAWG training set and the independent validation data sets, a mixture of different human genome builds, alignment algorithms and SNV calling algorithms were used for the validation data sets. We did not attempt to recall the SNVs, but did lift the genome coordinates of samples that had been aligned to other genome builds to hg19 by CrossMap (Version 0.2.5).

*Machine learning procedure - Random Forest*

For each of the 24 cancer types selected from the PCAWG sample set, we first used Random Forest model to train classifiers for each cancer type on each of the feature categories described in the above section. The data sets were z-score normalized across the samples before training. We used nested cross validation to train and test the performance of the classifiers. In the outer loop, the data set was divided into four folds and each fold was later used as an independent testing set. In the inner loop, the training portion of the data set was split into three folds and each fold was used as validation data set to fine-tune the hyperparameters. In the inner loop, we first used a chi squared test to filter out non-informative (V coefficient equals to 0) features. Then we tuned two hyperparameters for the Random Forest model to achieve the highest cross-validation F1 score. The two hyperparameters were the sample size for positive versus negative classes and the number of trees. We used the default R randomForest package parameter settings to sample the square root of the number of features at each split of the tree. The code was written in R (version 3.3.0). The main packages used were MLR (version 2.11) and randomForest (4.6-12) in training the model.

*Machine learning procedure - Neural Network Neural Network*

We ultimately used a fully-connected, feed-forward neural network for the classification of the 24 cancer types based on SNV type and mutational distribution alone. The network had a softmax output, which can be interpreted as a probability distribution of the 24 types. The predicted tumour type was selected by taking the type with the greatest softmax probability.

We used a Bayesian optimization approach to select hyperparameters^5^. Prior to training, data from PCAWG was split into training, validation and test sets 10 times to create 10 different partitions over the full dataset. For each of the 10 partitions, hyperparameters were selected by optimizing performance on the validation data for that partition. We used the `gp_minimize` function from the scikit-optimize 0.5.2 python library^6^ to select the following hyperparameters: learning rate for Adam, L2-regularization penalty (otherwise known as weight decay), dropout rate^7^, the number of hidden layers, the number of neurons per hidden layer, and activation function. Each model was trained using Adam^8^ with a batch-size of 32 for 50 epochs. All hyperparameters of Adam other than learning rate were set to the default values specified in the original paper^4^. Bias values were initialized as 0, and all other network weights were initialized using a glorot uniform distribution^9^. The model was evaluated with 200 hyperparameter combinations (i.e., 200 calls to `gp_minimize` were made). Briefly, `gp_minimize` approximates a function of model performance based on the hyperparameters with a Guassian Process. For each function call to `gp_minimize`, the performance on the current set of hyperparameters is evaluated by training the neural network, and assessing accuracy on the validation set. Based upon this accuracy, the Guassian Process is updated, and a new set of hyperparameters is chosen by optimizing an acquisition function. We used expected improvement as the acquisition function. After hyperparameter optimization, model performance was assessed independently on the corresponding test set for that split. Supplementary Table 6 describes the settings for each of the folds for these hyperparameters.

In order to compare the accuracy of these models with models trained on different feature sets, the procedure above was repeated using driver genes/pathways as input, and again by appending the driver genes/pathways features to the SNV features used above. The final hyperparameter values and model accuracies for each of the trained models is described in Supplementary Table 7. Each model was implemented and trained in Tensorflow 1.10.0^10^ and Keras 2.1.5^11^. All code was written in Python 3.6.

*Definitions of Accuracy Metrics*

To measure the performance of the classifiers, we use the conventional definitions of recall, precision, F1 score and accuracy. In the descriptions below, we use the abbreviations TP (true positive), TN (true negative), FP (false positive), and FN (false negative) to describe correct and incorrect assignments of an unknown tumour to a predicted type, as described by this confusion matrix:

| *Is the unknown sample a member of a particular histopathological type?* | **Predicted Yes** | **Predicted No** |
| --- | --- | --- |
| **Actually Yes** | *TP* | *FN* |
| **Actually No** | *FP* | *TN* |

***Recall****:* The proportion of samples of a particular histopathological type that are correctly assigned to that type:

Recall = TP/(TP+FN)

***Precision*:** The proportion of samples assigned to a particular type that are truly that type:

Precision = TP/(TP+FP)

***F1 Score:*** The harmonic mean of recall and precision:

F1 = 2(recall*precision)/(recall+precision)

**Accuracy:** The proportion of correct assignments. We use this metric only when summarising the performance of the classifier across all 24 tumour types:

Accuracy	= (TP+TN)/(TP+FP+TN+FN)

= Correct Assignments/Total Samples

*Literature Cited*

1. David Croft, Gavin O’Kelly, Guanming Wu, Robin Haw, Marc Gillespie, Lisa Matthews, Michael Caudy, Phani Garapati, Gopal Gopinath, Bijay Jassal, Steven Jupe, Irina Kalatskaya, Shahana Mahajan, Bruce May, Nelson Ndegwa, Esther Schmidt, Veronica Shamovsky, Christina Yung, Ewan Birney, Henning Hermjakob, Peter D’Eustachio, and Lincoln Stein. Reactome: a database of reactions, pathways and biological processes. Nucleic Acids Research. 39:D691-D697. (2011).
2. Alexandrov *et al.* The Repertoire of Mutational Signatures in Human Cancer. BioRixv preprint DOI <https://doi.org/10.1101/322859>
3. Aung, K. L., Fischer, S. E., Denroche, R. E., Jang, G.-H., Dodd, A., Creighton, S., et al. Genomics-Driven Precision Medicine for Advanced Pancreatic Cancer: Early Results from the COMPASS Trial. Clinical Cancer Research, 24(6), 1344–1354. (2018).
4. Priestley P *et al.* Pan-cancer whole genome analyses of metastatic solid tumours. BioRixv preprint DOI <https://doi.org/10.1101/415133>.
5. Snoek, J., Larochelle, H. & Adams, R. P. *Practical Bayesian Optimization of Machine Learning Algorithms*.
6. Head, T. *et al.* scikit-optimize/scikit-optimize: v0.5.2. (2018). doi:10.5281/ZENODO.1207017
7. Srivastava, N., Hinton, G., Krizhevsky, A. & Salakhutdinov, R. *Dropout: A Simple Way to Prevent Neural Networks from Overfitting*. *Journal of Machine Learning Research* **15,** (2014).
8. Kingma, D. P. & Ba, J. Adam: A Method for Stochastic Optimization. (2014).
9. Glorot, X. & Bengio, Y. *Understanding the difficulty of training deep feedforward neural networks*.
10. Abadi, M. *et al.* *TensorFlow: Large-Scale Machine Learning on Heterogeneous Distributed Systems*.
11. Chollet, F. Keras. *https://keras.io* (2015). Available at: https://keras.io/. (Accessed: 1st June 2018)
