## Supplementary Figure Captions for "A deep learning system can accurately classify primary and metastatic cancers based on patterns of passenger mutations"

**Supplementary Figure 1:** Radar plots showing the accuracy of classifier models constructed using single feature categories across all 24 tumor types. The distance from the center of the plot indicates the average F1 score for each model during nested cross-validation of the test data set.

**Supplementary Figure 2:** .Heatmap displaying the accuracy of the merged classifier using a held out portion of the PCAWG data set for evaluation. Each row corresponds to the true tumor type; Columns correspond to the predictions emitted by each of the classifiers. Cells are labeled with the proportion of tumors of a particular type that were called by each type-specific classifier. The recall and precision of each classifier is shown in the color bars at the top and left sides of the matrix.

**Supplementary Figure 3:** Violin charts demonstrating the distribution of the number of SNVs in the PCAWG and validation data sets. Note that we have paired the validation set of pediatric gliomas with the PCAWG juvenile piloastrocytoma data set.
