## Supplementary Figure 1 for "A deep learning system can accurately classify primary and metastatic cancers based on patterns of passenger mutations"

Supplementary Figure 1: Radar plots for all tumor types

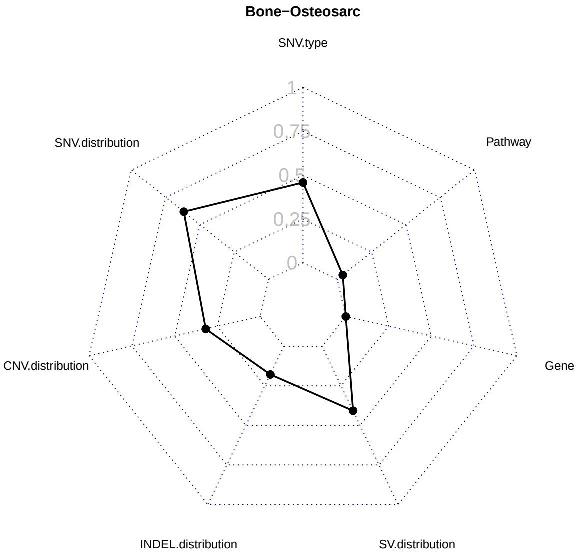
Bone−Osteosarc
SNV.type
Pathway
Gene
SV.distribution
INDEL.distribution
CNV.distribution
SNV.distribution
1
0.75
0.5
0.25
0

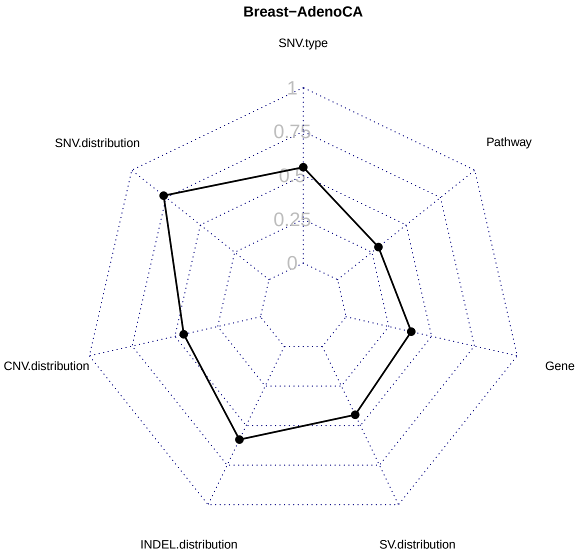
Breast−AdenoCA
SNV.type
Pathway
Gene
SV.distribution
INDEL.distribution
CNV.distribution
SNV.distribution
1
0.75
0.5
0.25
0

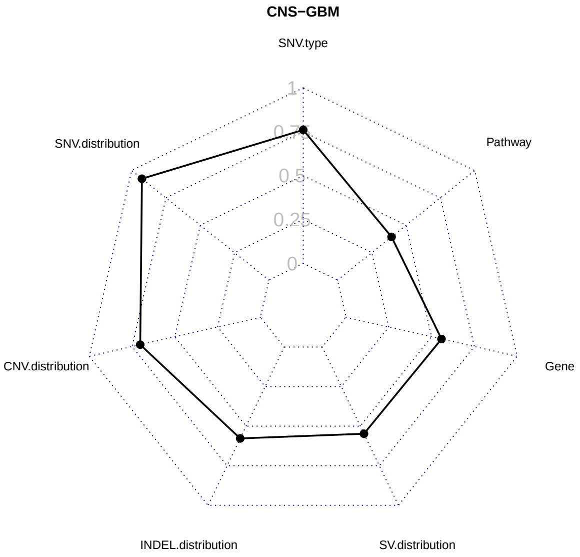
CNS−GBM
SNV.type
Pathway
Gene
SV.distribution
INDEL.distribution
CNV.distribution
SNV.distribution
1
0.5
0.25
0

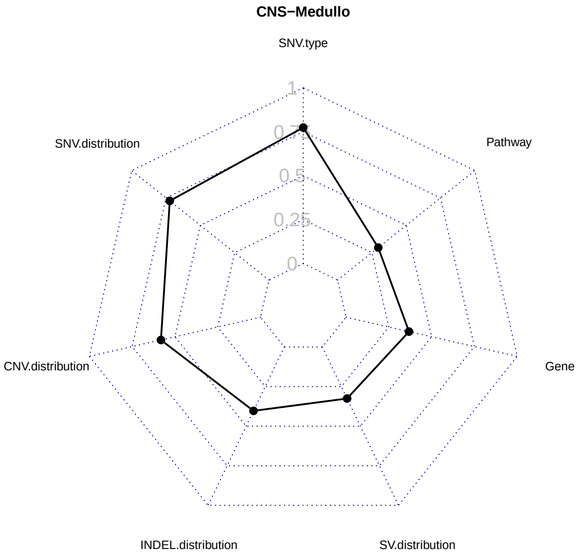
CNS−Medullo
SNV.type
Pathway
Gene
SV.distribution
INDEL.distribution
CNV.distribution
SNV.distribution
1
0.5
0.25
0

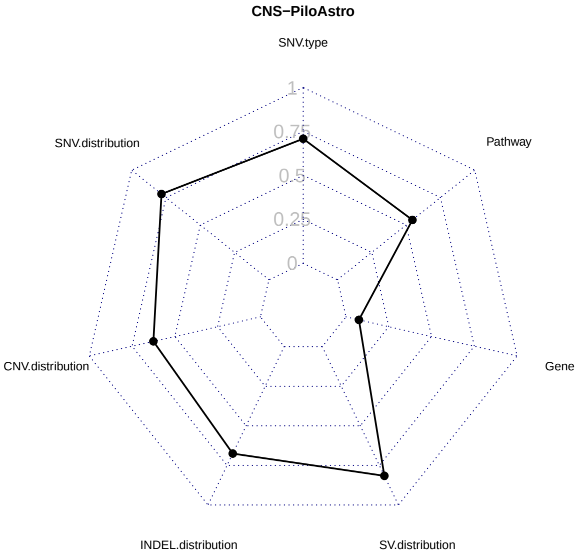
CNS−PiloAstro
SNV.type
Pathway
Gene
SV.distribution
INDEL.distribution
CNV.distribution
SNV.distribution
1
0.75
0.5
0.25
0

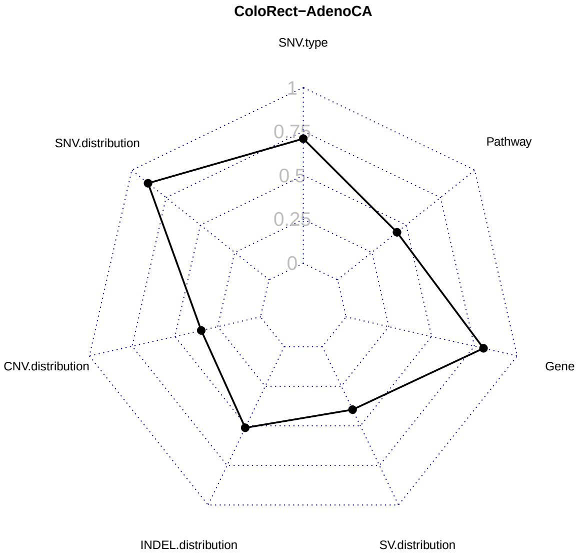
ColoRect−AdenoCA
SNV.type
Pathway
Gene
SV.distribution
INDEL.distribution
CNV.distribution
SNV.distribution
1
0.75
0.5
0.25
0

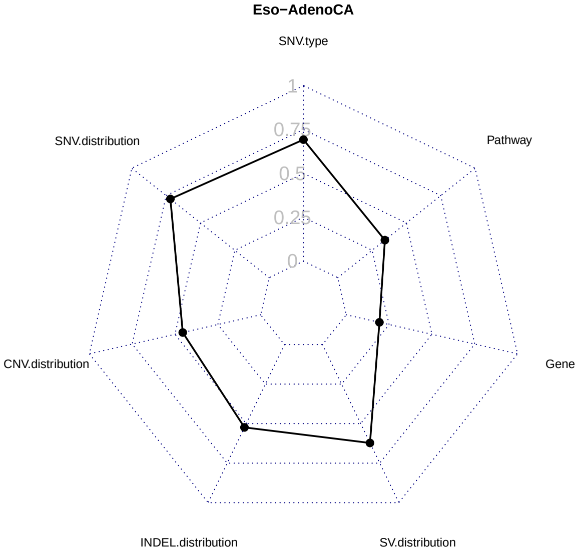
Eso−AdenoCA
SNV.type
Pathway
Gene
SV.distribution
INDEL.distribution
CNV.distribution
SNV.distribution
1
0.75
0.5
0.25
0

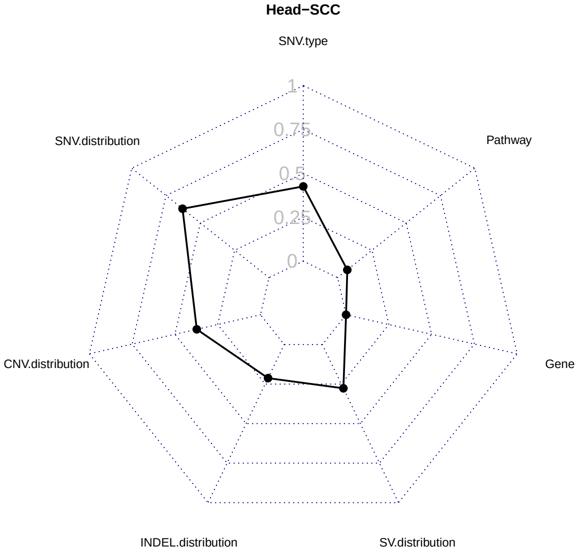
Head−SCC
SNV.type
Pathway
Gene
SV.distribution
INDEL.distribution
CNV.distribution
SNV.distribution
1
0.75
0.5
0.25
0

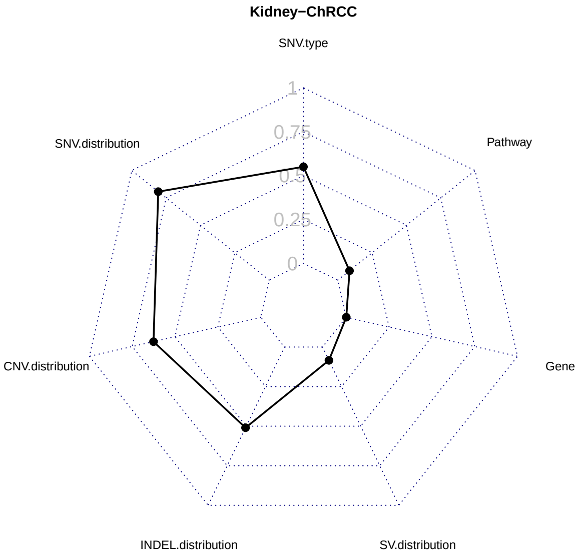
Kidney−ChRCC
SNV.type
Pathway
Gene
SV.distribution
INDEL.distribution
CNV.distribution
SNV.distribution
1
0.75
0.5
0.25
0

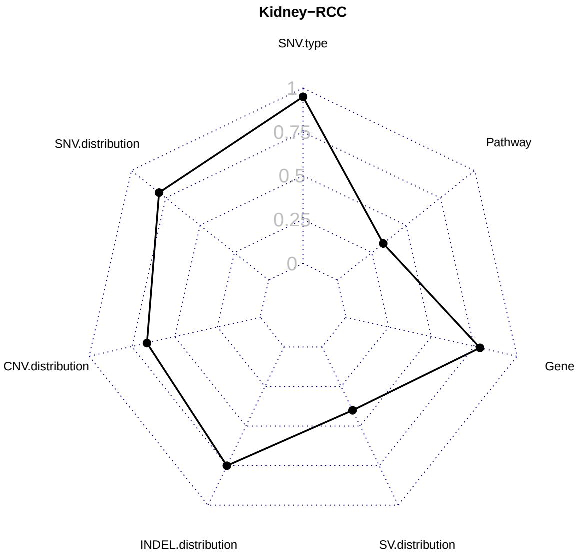
Kidney−RCC
SNV.type
Pathway
Gene
SV.distribution
INDEL.distribution
CNV.distribution
SNV.distribution
1
0.75
0.5
0.25
0

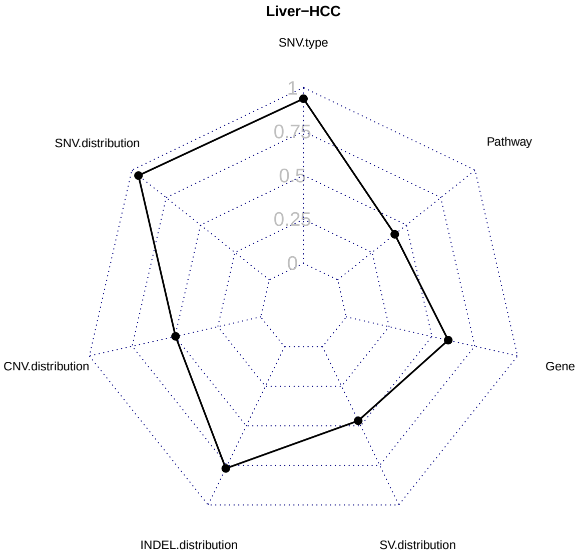
Liver−HCC
SNV.type
Pathway
Gene
SV.distribution
INDEL.distribution
CNV.distribution
SNV.distribution
1
0.75
0.5
0.25
0

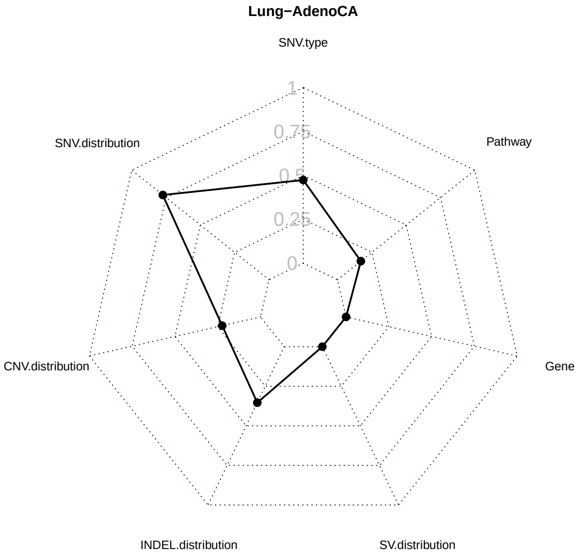
Lung−AdenoCA
SNV.type
Pathway
Gene
SV.distribution
INDEL.distribution
CNV.distribution
SNV.distribution
1
0.75
0.5
0.25
0

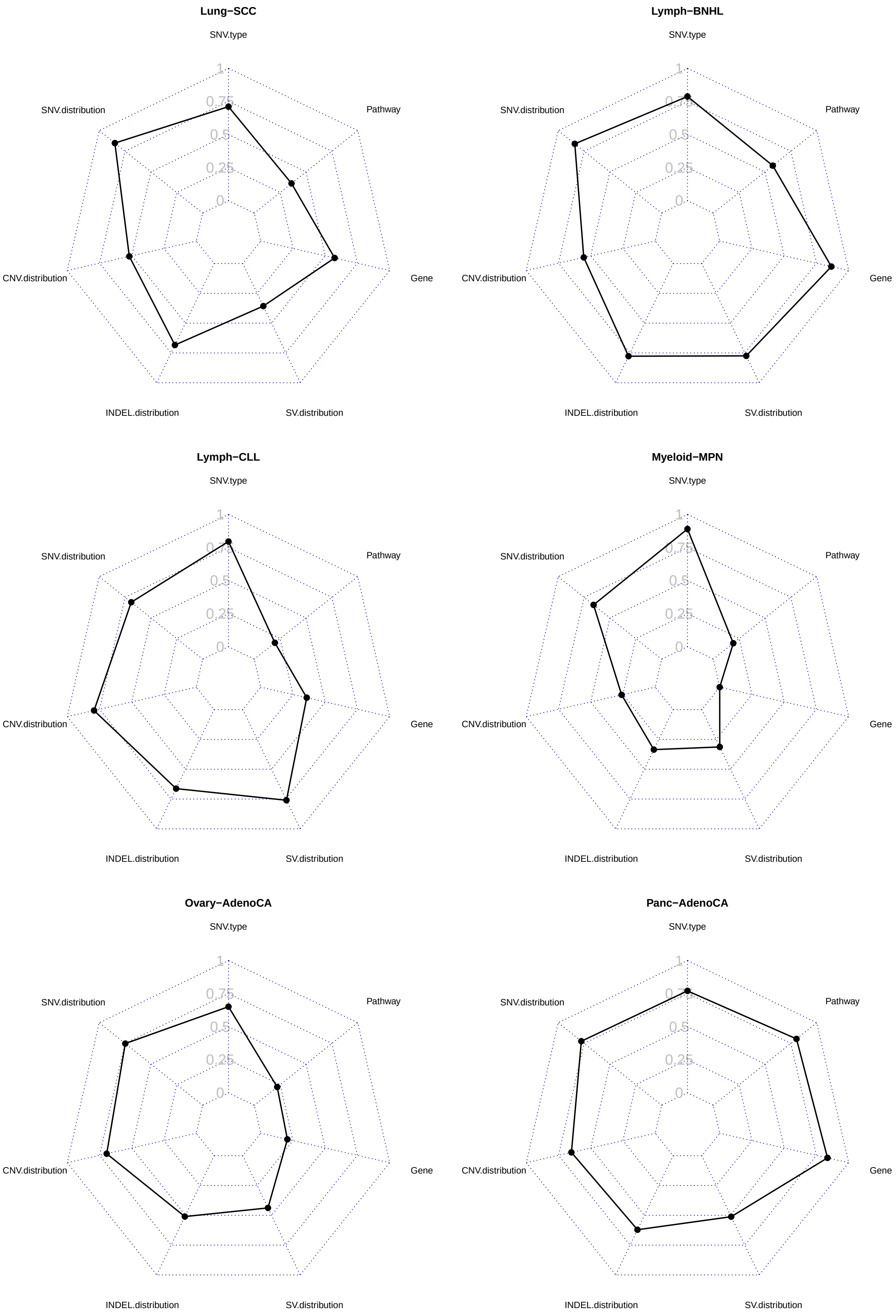
Lung−SCC
SNV.type
Pathway
Gene
SV.distribution
INDEL.distribution
CNV.distribution
SNV.distribution
1
0.75
0.5
0.25
0
Lymph−BNHL
SNV.type
Pathway
Gene
SV.distribution
INDEL.distribution
CNV.distribution
SNV.distribution
Lymph−CLL
SNV.type
Pathway
Gene
SV.distribution
INDEL.distribution
CNV.distribution
SNV.distribution
Myeloid−MPN
SNV.type
Pathway
Gene
SV.distribution
INDEL.distribution
CNV.distribution
SNV.distribution
Ovary−AdenoCA
SNV.type
Pathway
Gene
SV.distribution
INDEL.distribution
CNV.distribution
SNV.distribution
Panc−AdenoCA
SNV.type
Pathway
Gene
SV.distribution
INDEL.distribution
CNV.distribution
SNV.distribution

**Panc−Endocrine**

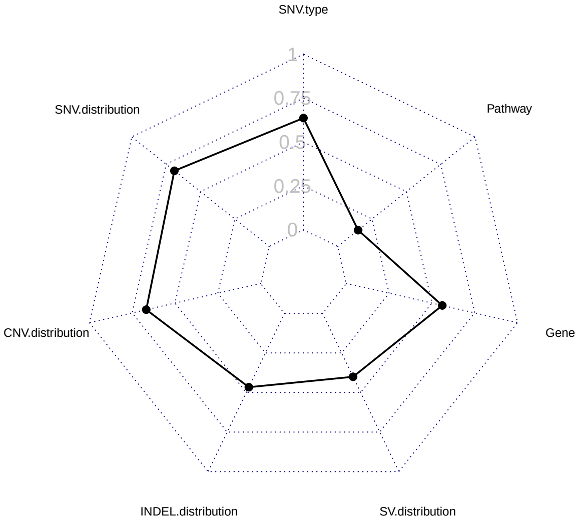


**Prost−AdenoCA**

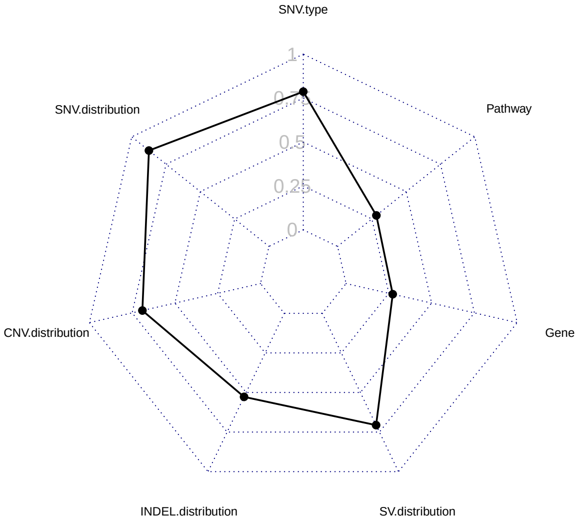


**Skin−Melanoma**

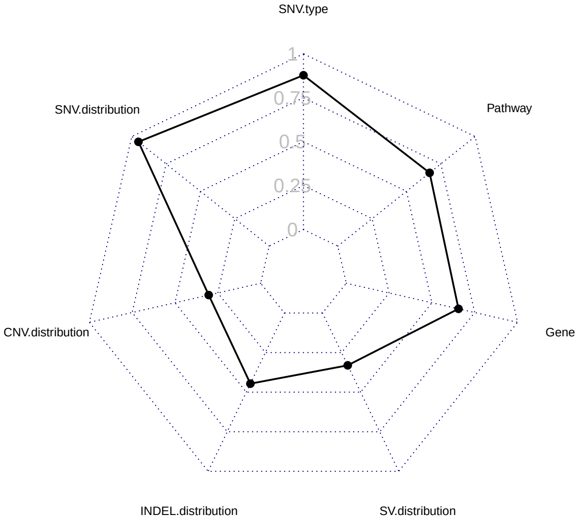


**Stomach−AdenoCA**

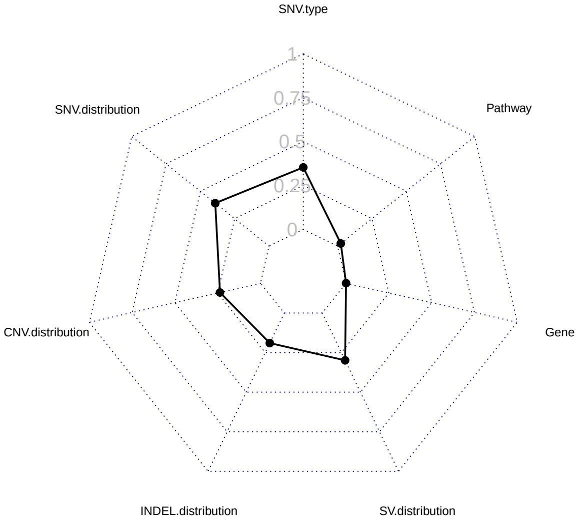


**Thy−AdenoCA**

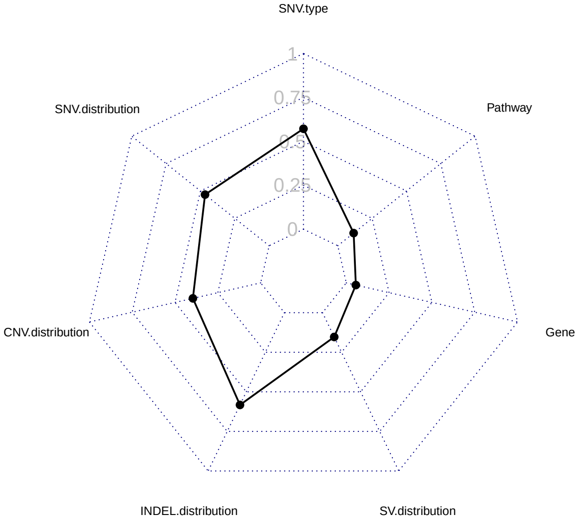


**Uterus−AdenoCA**

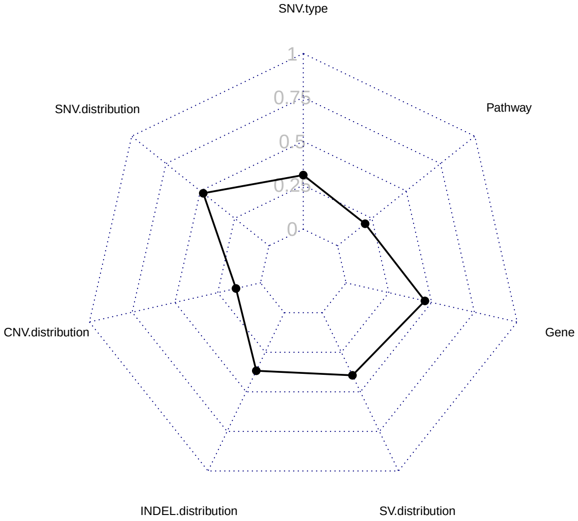
