## Supplementary figures and images for "A deep learning system can accurately classify primary and metastatic cancers based on patterns of passenger mutations"

### Supplementary Figure 2

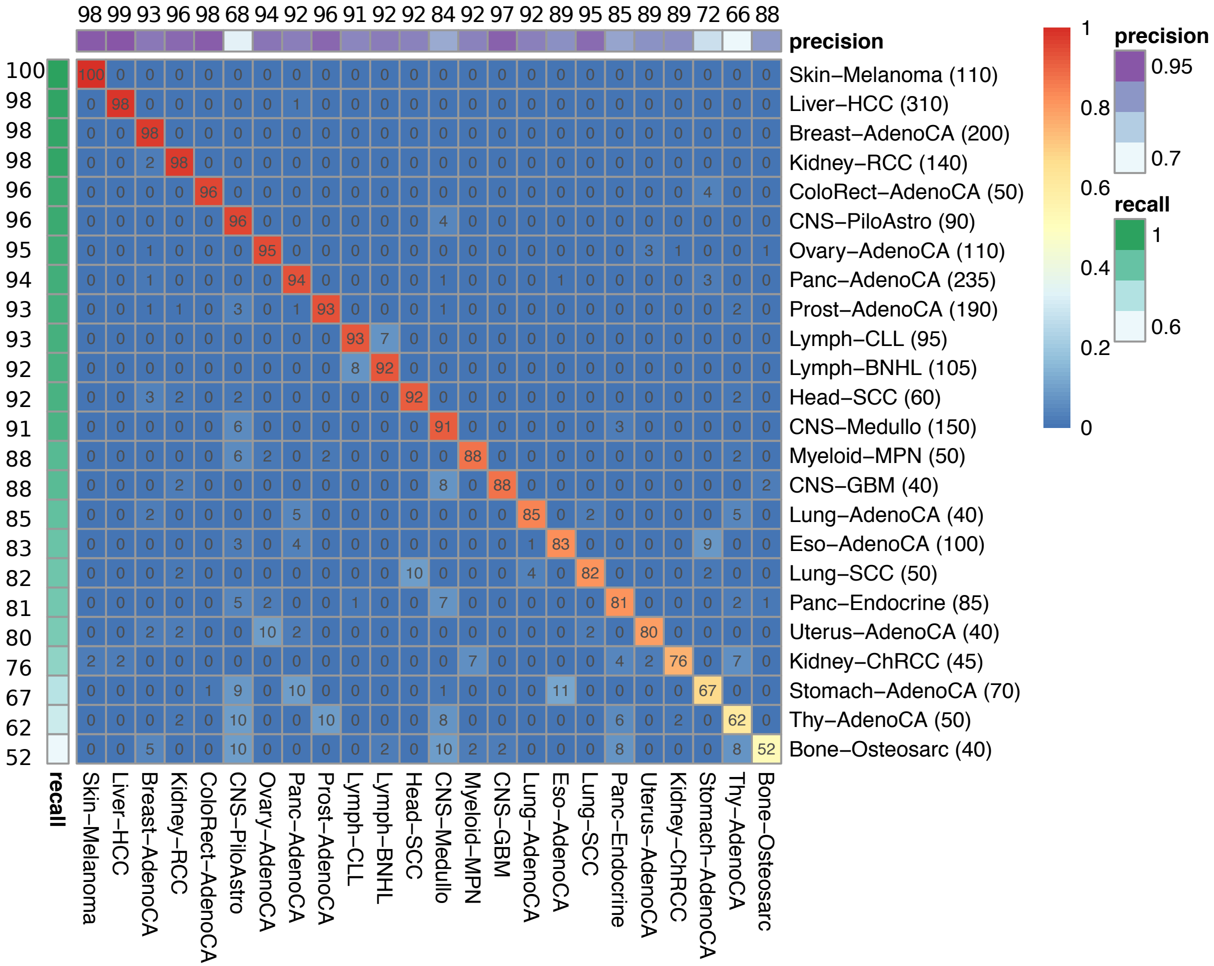

### Supplementary Figure 3

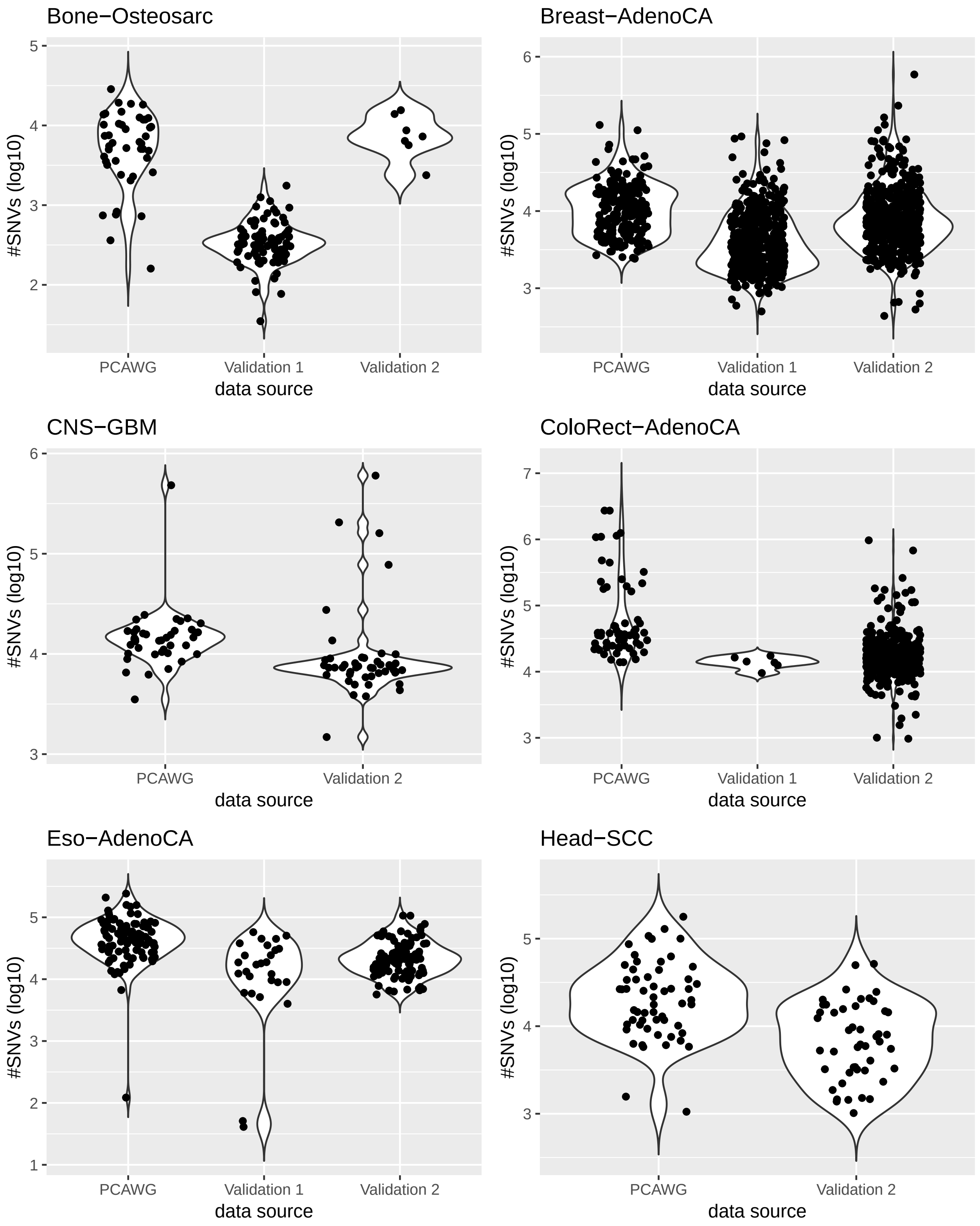

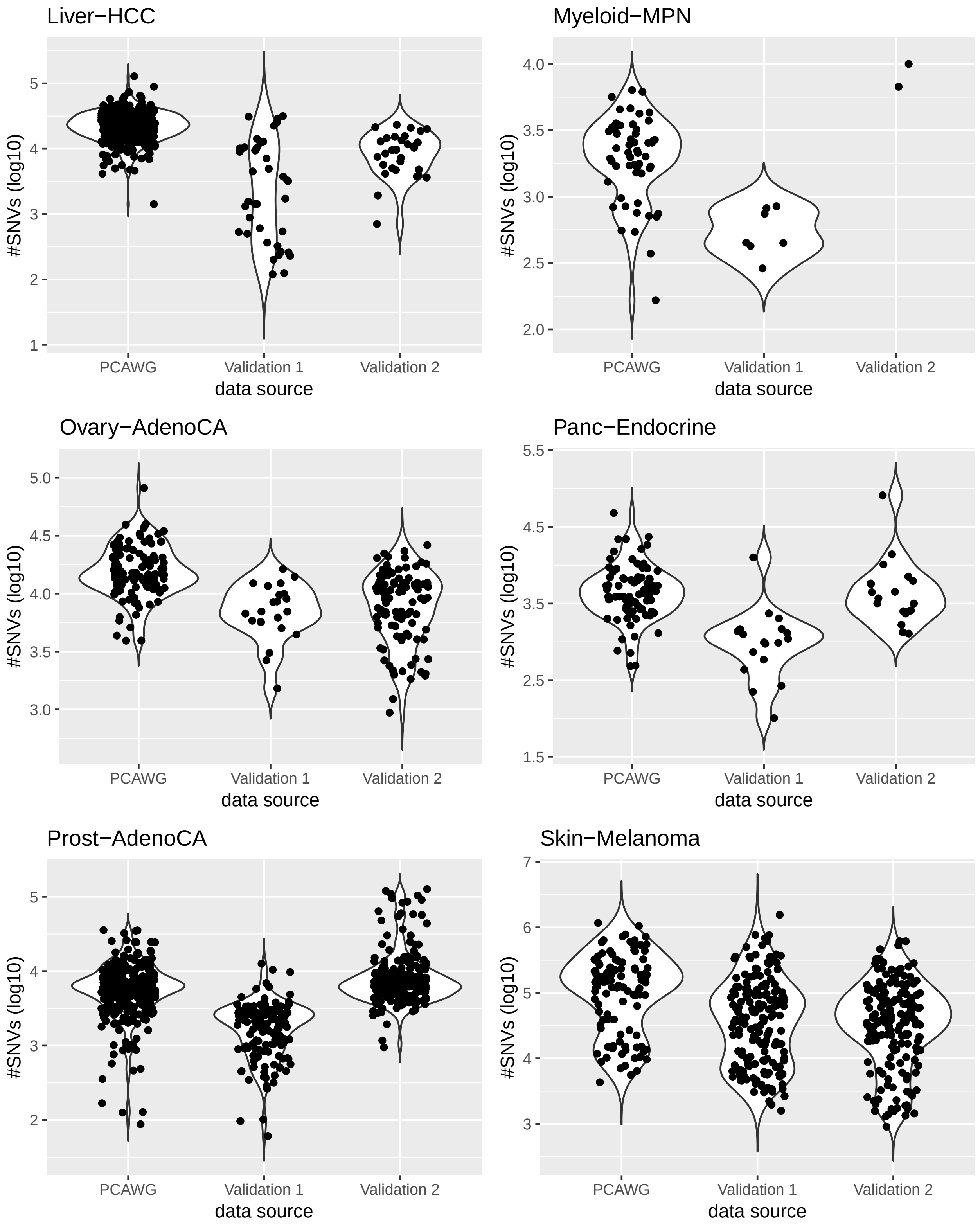

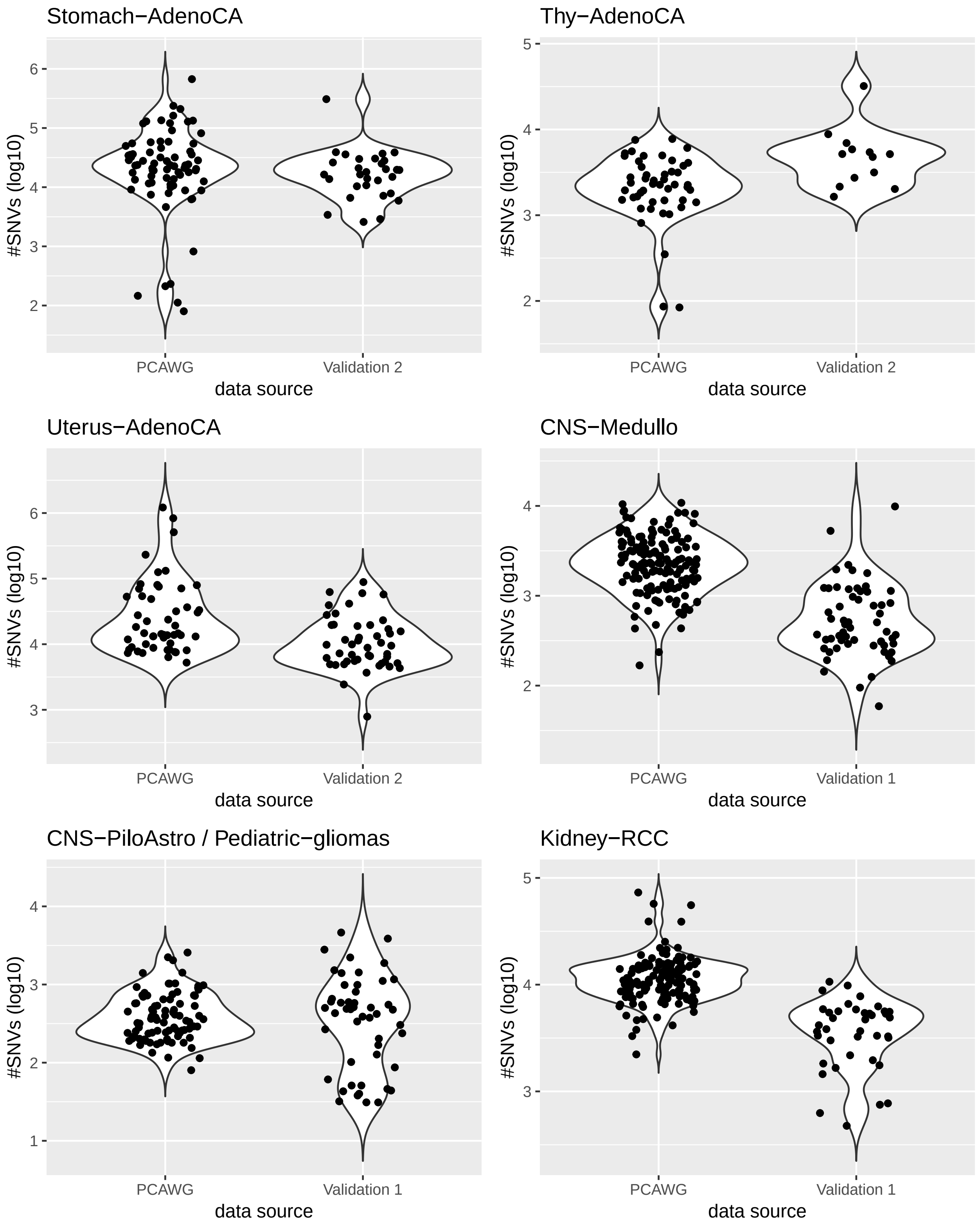

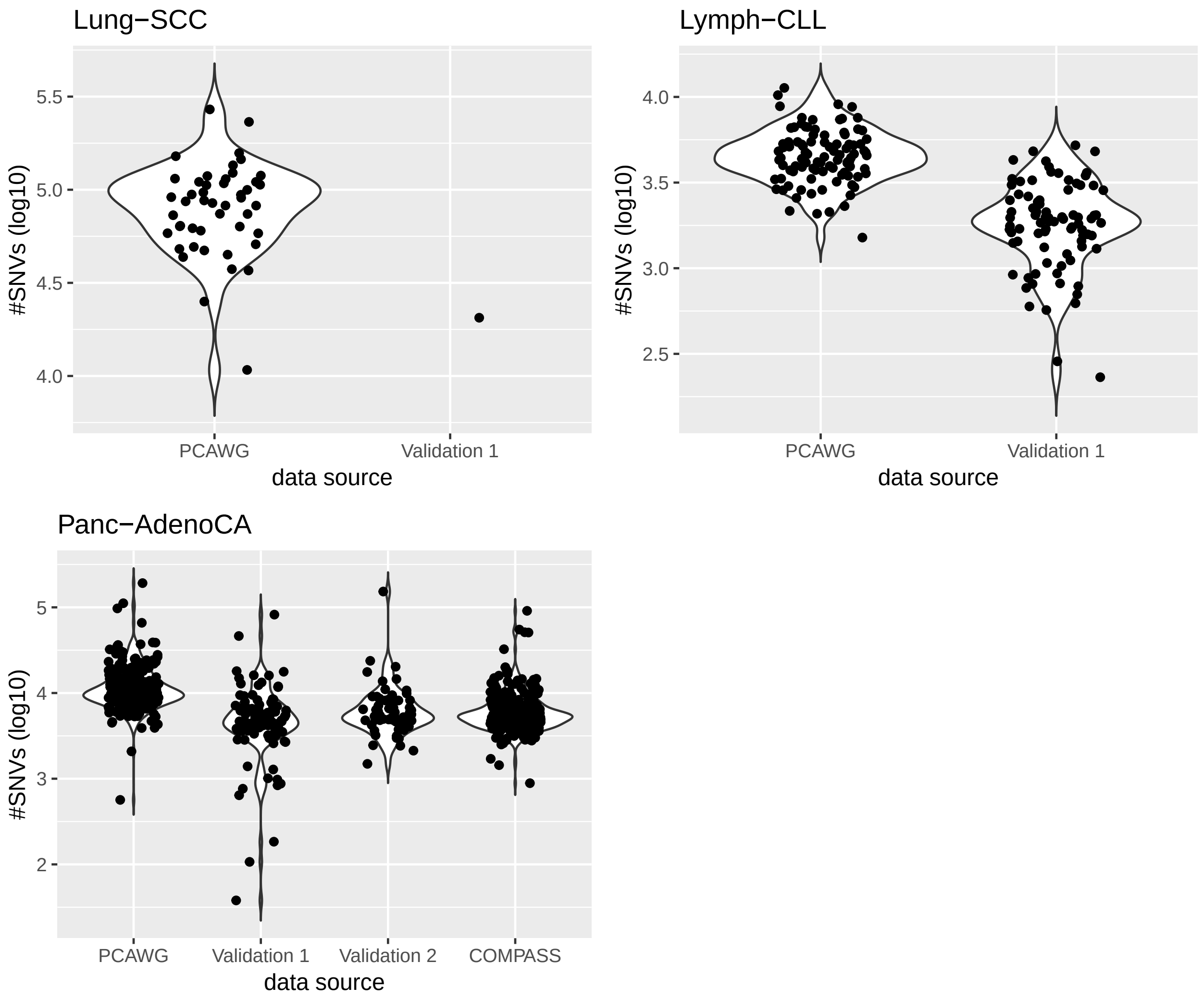
